# The antiviral mechanism of MxA revealed by a molecular docking-based structural model of the complex of MxA and influenza A virus NP/RNPs

**DOI:** 10.1101/219915

**Authors:** Yeping Sun

## Abstract

MxA, a member of the dynamin superfamily, is a potent host restriction factor for influenza A virus replication. The viral nucleoprotein (NP) of influenza A virus has been suggested to be a target of MxA. However, the molecular details of the interactions between NP and MxA remain unknown. In the present study, loop L4 of MxA, which was shown to target viral component and is missing in the crystal structure of MxA was modeled on a dimer of the MxA. The dimer was then docked onto NP. The resulting NP-MxA dimer complex is in agreement with previous studies, as the interface contains many residues that were previously identified to be associated with MxA resistance. And the model was the used to construct NP-MxA ring. We also constructed a structural model of the complex of influenza virus NP and polymerase, and the NP residues that interact with MxA in the NP-MxA model partly overlap with those in the NP-polymerase model, so MxA may competitively inhibit the binding of NP and the polymerase and disrupt the assembly of ribonucleoprotein (RNP) complex. These results of this work represent a putative molecular mechanisms for MxA antiviral activity.

## Introduction

Avian influenza viruses (AIVs) are zoonotic pathogens that mainly circulate in birds. They occasionally cross the species barrier and infect mammals and humans(1). All the four major human influenza pandemics since the beginning of the last century have been genetically related to AIVs. For example, the 1918 pandemic virus originated shortly before 1918 when a human H1 virus acquired avian N1 neuraminidase and internal protein genes (1). The H2N2 pandemic virus of 1957 was generated by reassortment of the previously circulating human H1N1 virus and an avian H2N2 virus that contributed the PB1, HA, and NA genes to the pandemic strain (2, 3). The 1968 H3N2 “Hong Kong” pandemic was caused by a reassortment event which resulted in a circulating human H2N2 virus acquiring avian HA (H3 subtype) and PB1 gene segments (4). Also in case of 2009 H1N1 swine-origin pandemic, reassortment produced the pandemic virus which processed two avian fragments (PB2 and PA) (4, 5). More recently, avian influenza A viruses of the H5N1 and H7N9 subtypes have caused hundreds of cases of human infections. So far, sustained human-to-human transmission of these viruses has not been reported (6). In order to infect humans and establish a stable virus linage influenza A viruses of avian origin has to break the species barrier and acquire the capacity of efficient transmission in humans. To overcome the species barrier, several adaptive mechanisms to the new human hosts are required. Besides adaptations to host factors that facilitate infection, such as the adaptation of HA proteins to host receptors (7, 8), another important adaptive mechanism is the counteraction against cellular restriction factors that inhibit virus replication (9).

Human MxA protein, a dynamin-like GTPase, was amongst the first identified Interferon stimulated genes (ISGs) that restricts the replication of influenza A virus ((10-(13) and a wide range of other negative and positive strand RNA viruses (14, 15). It may thus function as an efficient barrier against zoonotic introduction of influenza A viruses into the human population. MxA is a cytoplasmic protein that associates with the smooth endoplasmic reticulum (ER) and the smooth ER-Golgi intermediate and sequester viral components upon virus infection (16, 17). Nucleoproteins (NPs) of influenza A virus have been identified as major target of MxA, and the interaction between NP and MxA has been studied in some detail. For example, a MxA construct targeted nucleus (nuclear MxA) can form complex with NP and inhibit the transcription step of the influenza virus genome, and the inhibitory activity of nuclear MxA was markedly neutralized by over-expression of the NP or an N-terminal, but not C-terminal region of the NP (18). Different strains of influenza A viruses showed different sensitivity to the antiviral activity of MxA. The pandemic 1918 H1N1 “Spanish flu” is insensitive to MxA, while the highly pathogenic avian H5N1 strain is sensitive (19). The different sensitivity of these influenza virus strains depend solely on their NP protein (20). By comparing the sequences of NP of highly pathogenic avian H5N1 strain, the pandemic 1918 H1N1 “Spanish flu”, and the 2009 H1N1 pandemic influenza A virus strain, a bunch of NP residues that determine the sensitivity of the virus stains to MxA are identified (9).

The MxA structure was recently determined (21, 22). It resembles that of other members of the dynamin-like large GTPase superfamily, consisting of an N-terminal GTPase domain (G domain) and a C-terminal stalk comprising an antiparallel four-helix bundle. These two structural domains are links by a bundle-signaling element (BSE) that is necessary to transfer structural changes during GTP binding and hydrolysis to the stalk. The crystal structures of the stalk and the close-to-full-length of MxA highlights the importance of self-assembly for the antiviral activity of MxA (21, 22). Thus, the stalk mediates dimerization of MxA via a highly conserved interface. Two further interface allow assembly of dimers into an oligomer. However, the crystal structure did not resolve the molecular basis for the long-standing problem how MxA mediates its antiviral specificity. This would require the atomic structure of the complex of MxA with its viral target components. Evolution analysis of the MxA coding regions of 24 primate species showed that the majority of positively selected sites were located in the MxA stalk, and the most striking enrichment of positively selected sites is an unstructured loop L4 at the tip of the stalk. This analysis revealed a long history of positive selection in L4 across primates, which suggested that the loop L4 was the genetic determinant of the antiviral specificity of Mx proteins (23).

Based on these studies, we made an attempt to construct a structural model of MxA to explain the interaction of MxA and the NP protein of influenza A virus.

## Results

### Modelling the missing loops in the crystal structures of MxA and H5 NP

Previous studies showed that influenza virus NPs are the target of MxA protein ((18, (19, (24). However, no atomic structure of the MxA-NP complex has been reported to date. This prevents a complete molecular understanding of the antiviral mechanisms of MxA. Here, we modelled the MxA-NP complex using molecular docking with the crystal structures of MxA and NP protein of influenza H5N1 virus.

In the published close-to-full-length MxA structure, some residues GTPase domain were disordered and not included in the model (residues 95-101, 128-130, 182-187, 265-266, 318-320, and 535-572). Most importantly, residues 535-572 comprising the L4 Loop were deleted in the crystallized construct (Figure 1A). In the H5N1 virus NP model (PDB code: 2Q06), residues 1-21 and 79-86 were missing (Figure 1B) (25). We first modelled the missing L4 fragment in MxA protein using the I-TASSER program (26, 27). I-TASSER provided five structural models of MxA (Figure S1 and S2). When constructing MxA dimers using these five MxA structural models, we found that in most dimers, the modelled L4 loop in one monomer clashed into the corresponding loop or the helical structures from the other monomer, or crash into the helix of the stalk in the other monomer (Figure S1). Only in one of these dimer, one MxA monomer did not clash with the opposing monomer (model 3, Figure 1C), so we use this dimer for the docking and construction of the higher-order MxA oligomer. As for H5N1 NP, we use the I-TASSER model with the best score (model1, Figure 1D) for docking.

**Figure 1.**
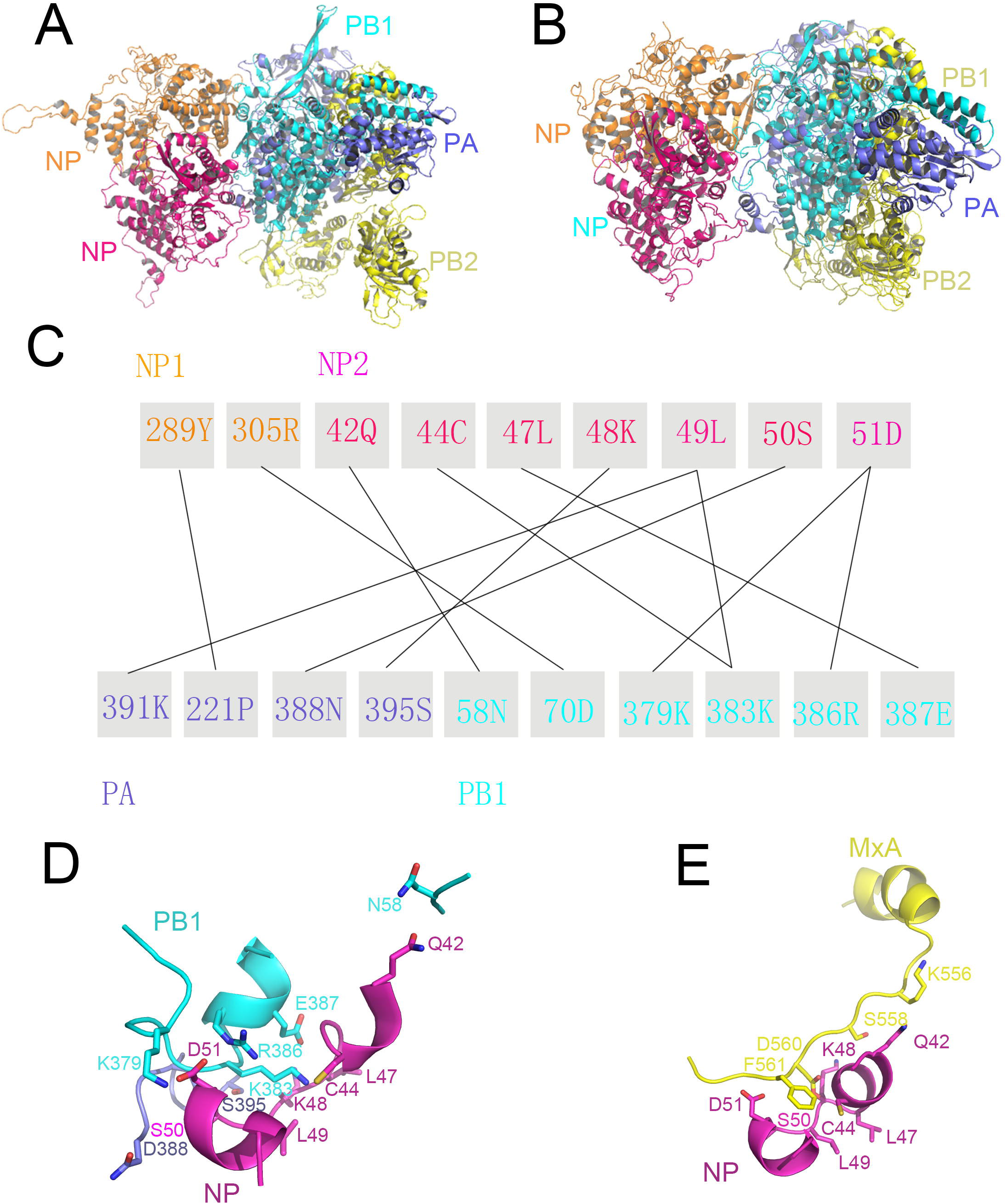
Mending the missing fragments in the crystal structures of MxA and H5 NP proteins. (A) The crystal structure of MxA. BSE: bundle signaling element. The missing residue 535-572, which may be involved in interact with viral targets, are indicated as dash line. (B) Crystal structure of H5 NP. The missing residue 1-21 and 535-572 are also indicated as dash line. (C) The structure of MxA with the missing fragments mended by I-TASSAR program. The mended residues 535-572 are shown as hotpink color. (D) The structure of NP with the missing fragments mended by ITASSAR server. The mended residues 1-21 and 535-572 are shown as hotpink color.

In model 3 of the MxA dimer, the two mended L4 loops formed a salient structure. They extend from α3^s^ region of the stalk, form a short helix, then tilt back with their C-terminal residues forming interactions with the α4^s^ region (Figure 2A). In the L4 loop, residue S568 forms interactions with M582 and Q586 in the α4^s^ region; A569 forms interactions with M582 and F578 in α4^s^; and T570 interacts with Q579 in α4^s^. Thus, residues K554 to A572 in the two L4 models form an extended and solvent-exposed interface that might interact with the viral targets (Figure 2B).

**Figure 2.**
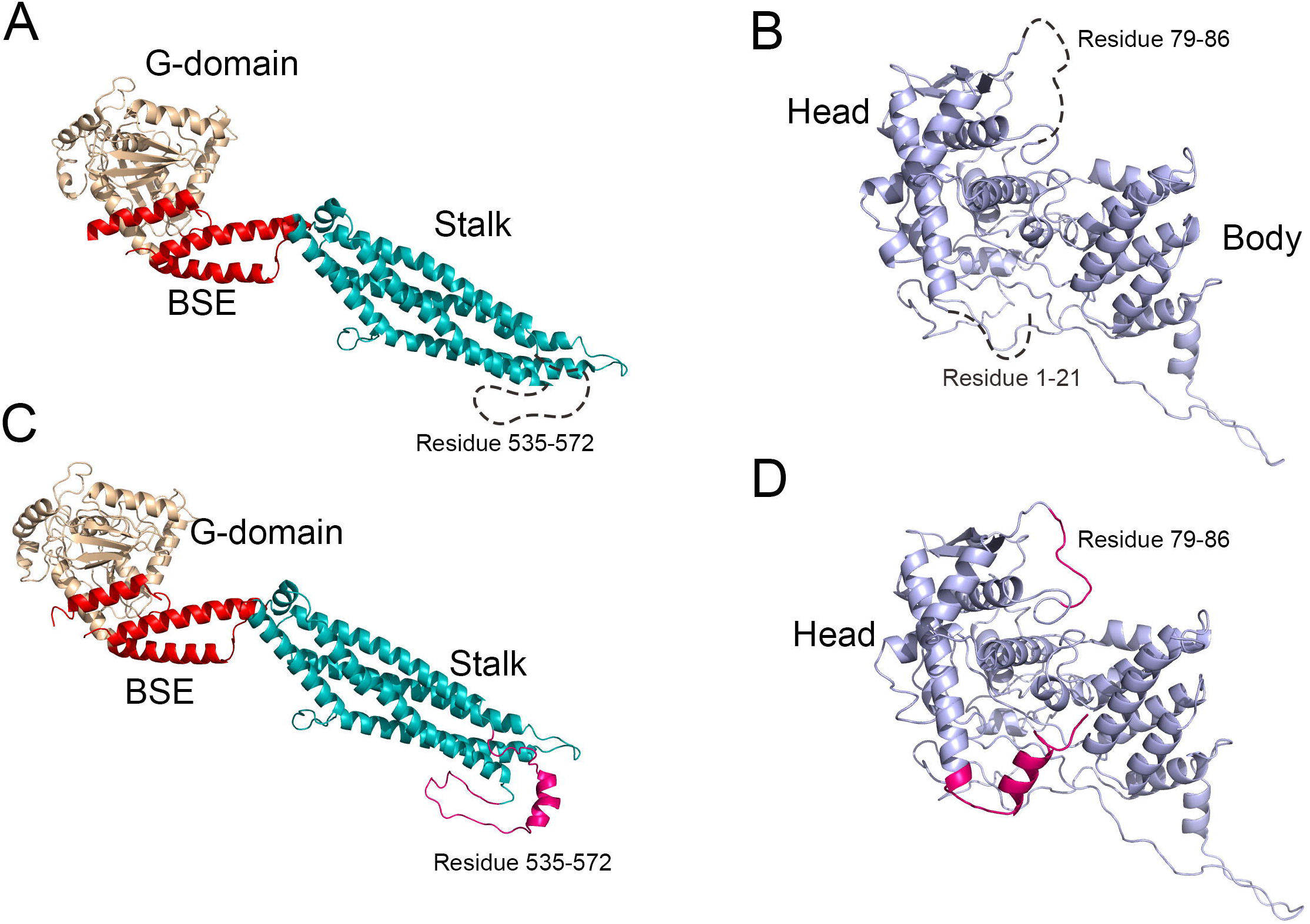
MxA dimer constructed with the MxA monomers mended by ITASSAR program. (A) The overall structure of the MxA dimer. (B) The L4 loops of the MxA dimer.

### Docking the MxA dimer to the NP of influenza H5N1 virus

Docking of the MxA dimer to NP was performed by Z-dock program (28). L4 of the two MxA monomers in the dimer were obvious exposed, so we set these residues as docking site. We did not set any restriction in the NP for docking. The Z-dock server yielded 10 models of NP-MxA dimer complexes. The interacting resides in the NP of the 10 complex models were listed in table S1. Recently, some mutations were reported to be associated with the escape of pandemic influenza A virus from human MxA restriction. This study showed that G16, Y100, L283, F313 were associated with MxA- resistance of the 1918 NP, while E53, R100, F313, Y289, R305, I316, T350, R351, and 5,T353 were relevant to resistance of the 2009 NP. We expect these residues localize at the interacting interface between MxA and NP (9). In these NPMxA dimer models, only one interface residue, R305 NP in model 2 and 5, is known to be associated with of NP resistance to MxA (table S1), whereas there are three such interacting interface residues, NP L283, Y289 and R305, in model 8 and 9. Because the interface in model 8 contains more interacting NP residues than in model 9, which suggests that the NP-MxA interaction can be stronger in model 8 than model 9, here we just select model 8 for further analysis.

**Table S1.**
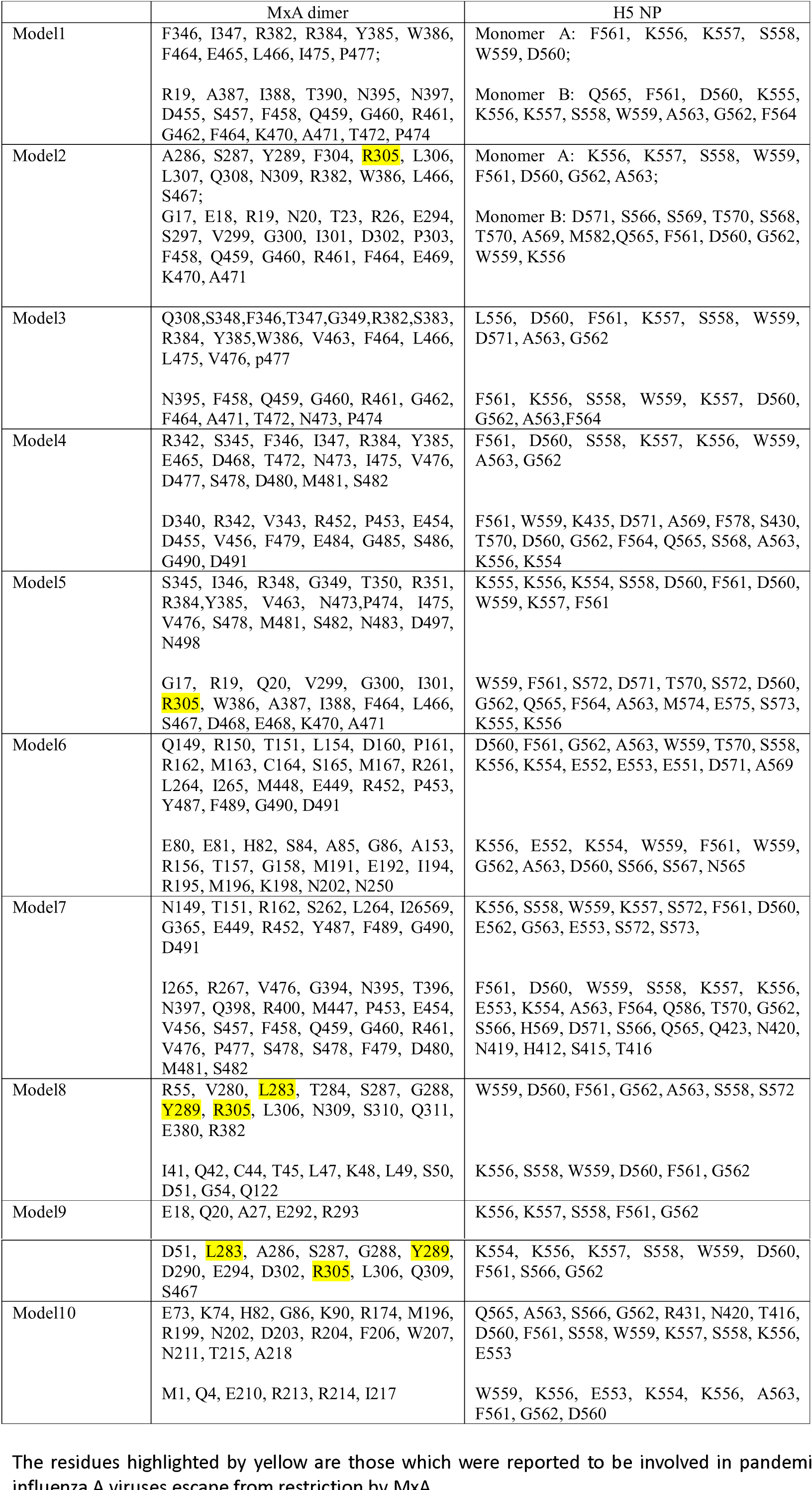
Residue contacts between the MxA dimer and H5 NP in the models of their complexes generated by Z-dock server.

In model 8, the two L4 loops bind to the head domain of NP, and the docking site is on the opposite side of the RNA binding groove (Figure 3A and 3B). The L4 loops extend along the surface of the NP head domain and form extensive interaction with the NP. There are totally 25 residues in NP that are involved in the interaction with MxA which comprise 9 pairs of hydrogen bonds, 1 pair of salt bridge, and numerous van der Waal interactions. Among the three residues linked to MxA-resistance, L283 of the NP locates to the center of the interaction interface (Figure 3C). It interacts with the aromatic ring of F561 in monomer 1 of the MxA dimer, but it also tends to interact with F561 in monomer 2 of the MxA dimer. This would explain how the L283P mutation in NP can compromise the antiviral activity of MxA: when the aliphatic side chain of Leu mutate, the interaction with the aromatic ring of F561 in MxA is lost. Y289 in NP, the second position associated with resistance to MxA, forms a hydrogen bond with D560 in monomer 1 of the MxA dimer, and it also form vdW interaction with F561 and G562. D350 in NP, the third residue position associated with MxA resistance, forms a salt bridge with monomer 1 of MxA, which is also important in stabilizing of the binding interface. In this way, the three residues in NP which are associated with MxA resistance contribute substantially to the binding of NP to monomer 1 of the MxA dimer. Besides, hydrogen bonds form between Q311 and R382 in the NP, and W559 and S558 in monomer 1 of the MxA dimer. On the other hand, monomer 2 of the MxA dimer mainly interacts with a continuous sequence, residue 41-54 in the NP. Interestingly, this sequence overlaps with one of the nucleus export signals (NES1, 24-49) of the NP (21).

**Figure 3.**
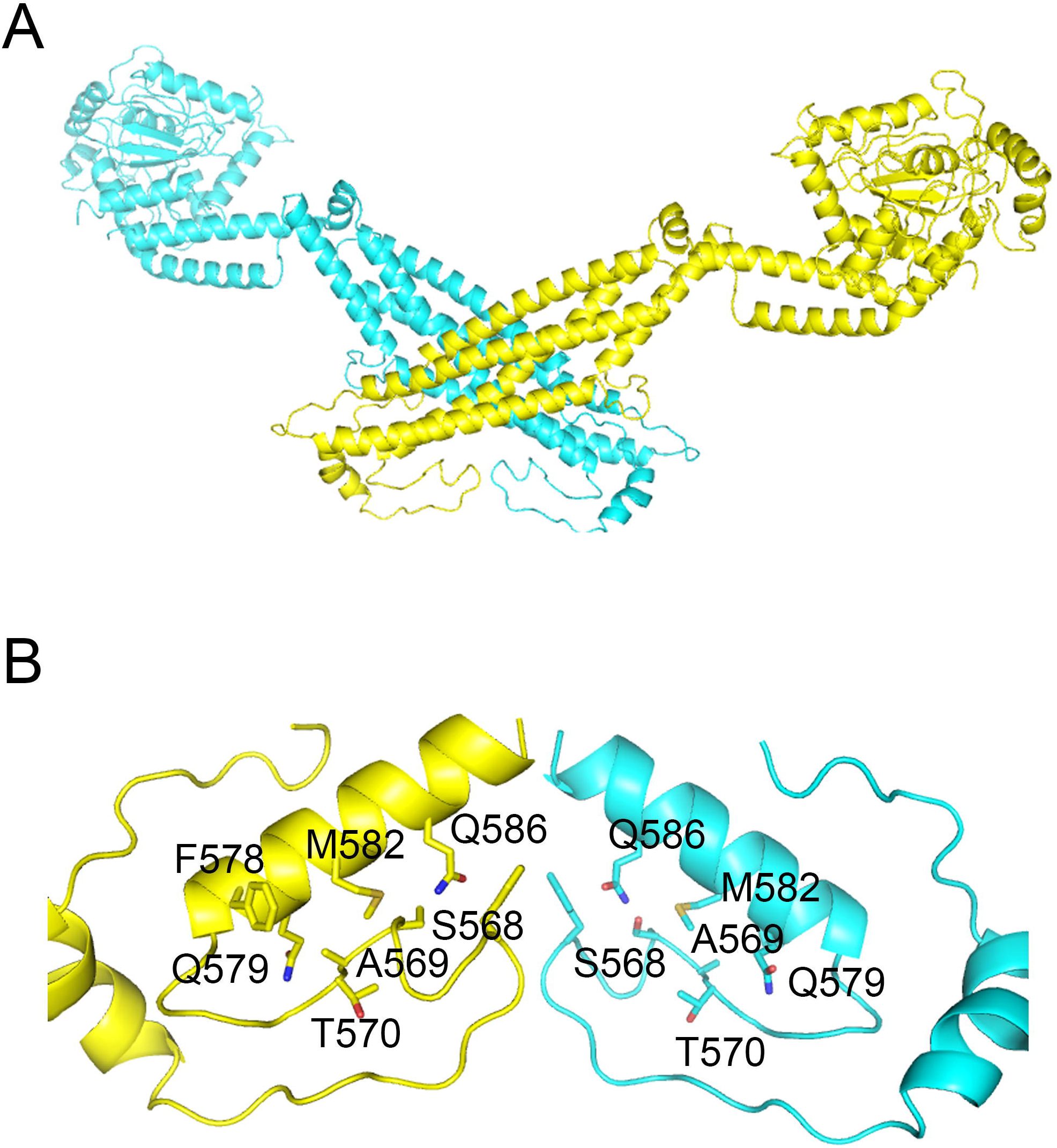
The complex structure (model 8) of NP-MxA dimer generated by Z-dock. (A) The cartoon representation of the overall structure of NP-MxA dimer. (B) The surface representation of the overall structure of NP-MxA dimer. NP is shown as coulombic surface (+/-5 kcal/ (mol e)) to highlight the electropositive RNA binding sites. (C) Comparison of the NP amino residues that are involved in the interface interactions between MxA dimer and NP. The virus strains in the comparison are: H5N1-NP, A/Hong_Kong/483/1997; 1998-NP: A/Brevig_Mission/1/1918; pH1N1-NP, A/California/04/2009; H7N9-NP, A/Guangdong/1/2013. (D) The interaction interface in the NP-MxA dimer complex. The interaction residues in MxA are shown as yellow; the interaction residues in NP are shown as hotpink. (E) The NP residues in the interaction interface in NP-MxA dimer (hotpink) and the NP residues that have been associated with MxA resistance (Yellow). The residues that are both involved in the interface interaction and associated with resistance to MxA are shown as blue.

When mapping the 25 NP interface residues in model 8 and the 13 NP residues that are associated with MxA resistance in NP structure of model 8 (Figure 3D), the relation between these two residue groups become clearer. Among the 13 NP residues that are associated with MxA resistance, three overlap with the NP interface in NP. Other five residues including E53, R100, F313, I316, T350 and R351 are near the interface, so mutations of them can also have certain effect on the interaction.

### The structure model of the NP-MxA ring and RNP-MxA ring complexes

Gao et al. previously presented a ring-like MxA oligomer model, which is composed of 16 MxA dimers (15). By aligning the Mx dimer to the proposed MxA oligomeric ring, we obtained an MxA ring with intact L4 loops. The solvent-exposed regions of L4 are still exposed and face to the cavity of the MxA ring (Figure 4A). By aligning our NP-MxA complex to the modeled MxA ring, we obtained a structural model of the ring-like MxA oligomer in complex with the NP (e.g. the NP-MxA ring). The cavity of MxA ring can accommodates five NP monomers, which are arranged at almost equal distance to each other in the cavity of the MxA ring (Figure 4B).

**Figure 4.**
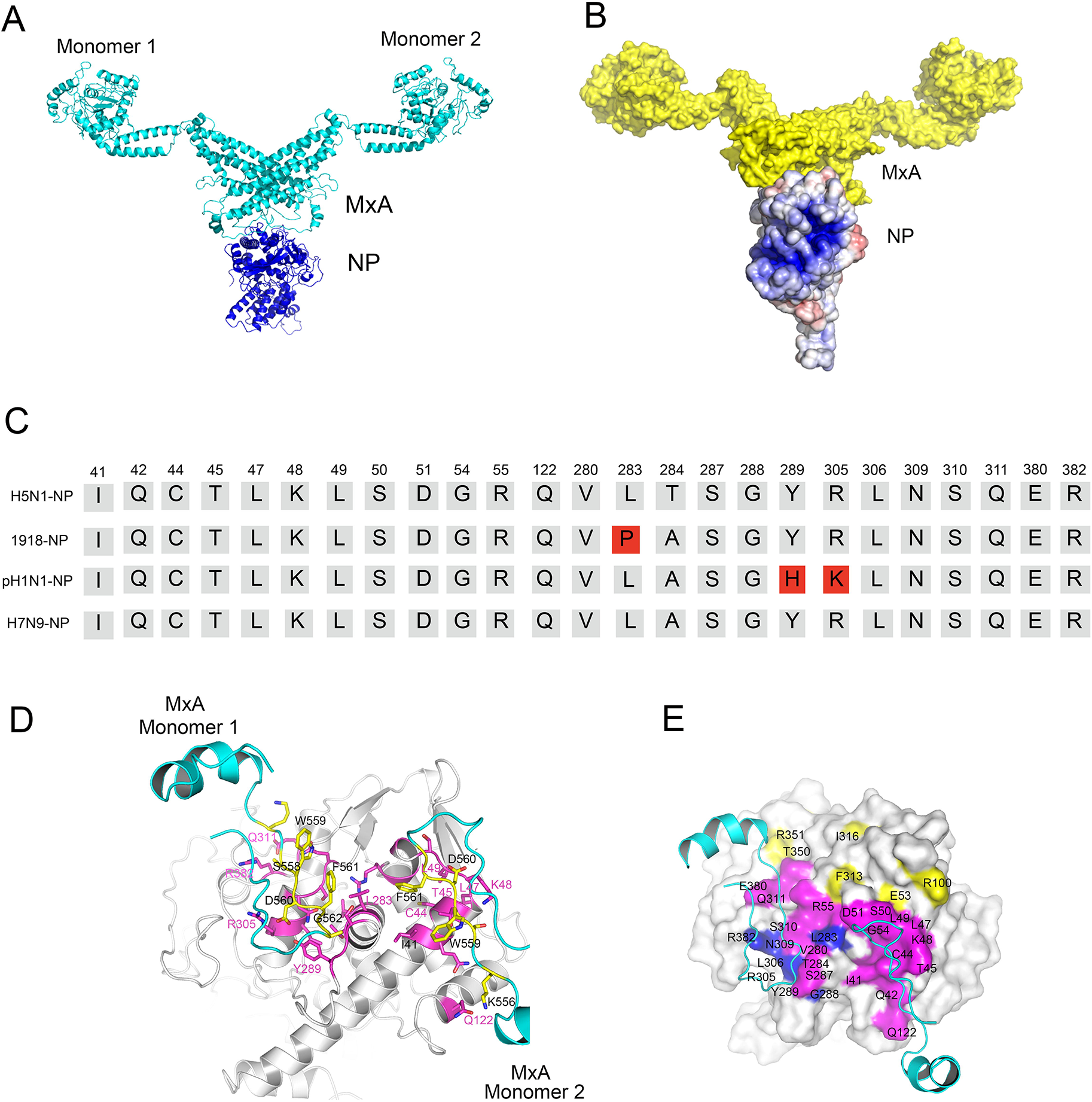
The structure models of the MxA ring with L4 loops and the NP-MxA ring complex. (A) The structure models of MxA ring with L4 loops was produced by align the 16 MxA dimers with modelled L4 loop to the MxA ring constructed with the MxA crystal structure in which the L4 loop is missing. (B) The structure model of NP-MxA ring complex was produced by aligning NP-MxA dimer to the structure models of the MxA ring.

Next, we examined how RNP could interaction with the MxA ring. Recent studies elucidated two structural models of the influenza vRNP complexes, derived either in its native form from purified virions or from cells transfected with plasmids that expressed the vRNP components, by using advanced electron microscopic analyses. These studies demonstrated that the vRNP internal region comprises an antiparallel double helix of vRNA–NP complexes (29, 30). The MxA interacting residues in NP we identified in the NP-MxA complex (figure 3C) localize along the surface of the helical RNP structure. By aligning the NP in the complex of NP-MxA ring (figure 4B) to each NP protomer in central helical filament of this RNP structure, a series of structural models of RNP-MxA complex could be obtained. However, there are obvious coordination clashes in all of these models, so the central helical filament of RNP may not be the binding site of MxA ring unless large conformational changes occur in the MxA ring or the MxA ring.

The complete RNP comprises the end loop, the central helical filament and the polymerase complex which comprises of PA, PB1 and PB2 proteins. To investigate whether the NP residues that interact with MxA in the NP-MxA complex model (figure 3A) also locate in the binding interface between the RNP central helical filament and the polymerase complex, we modelled NP-polymerase complex. We first modelled H5N1 polymerase complex structure by I-TASSER program (27), using the crystal structure of the bat influenza A virus polymerase (31) as the template (Figure S2A-S2D). Then we docked this H5N1 polymerase structural model (Figure S2D) into the electron density map of the RNP-bound polymerase (EMDB entry numbers: 2212) (30), together with two NP molecules which make contact with the polymerase. The resulting atomic model (Figure 5A) of the NP-polymerase complex were further refined by 2 nanosecond (ns) molecular dynamics flexible fitting (NAMD) (32). The Root Mean Square Deviation (RMSD) and CCC curves became satisfyingly converged at the end of the simulation (figure S4C and 3D). Compared with the initial structure, the final structural became obviously more compact and more closed fitted into the electron map (Figure S4A and 4B). The CCC value of this structure to the initial structure is 0.848). The last snapshot of the simulation (Figure 5B) was chosen as the final atomic model of NP-polymerase complex for further analysis.

**Figure 5.**
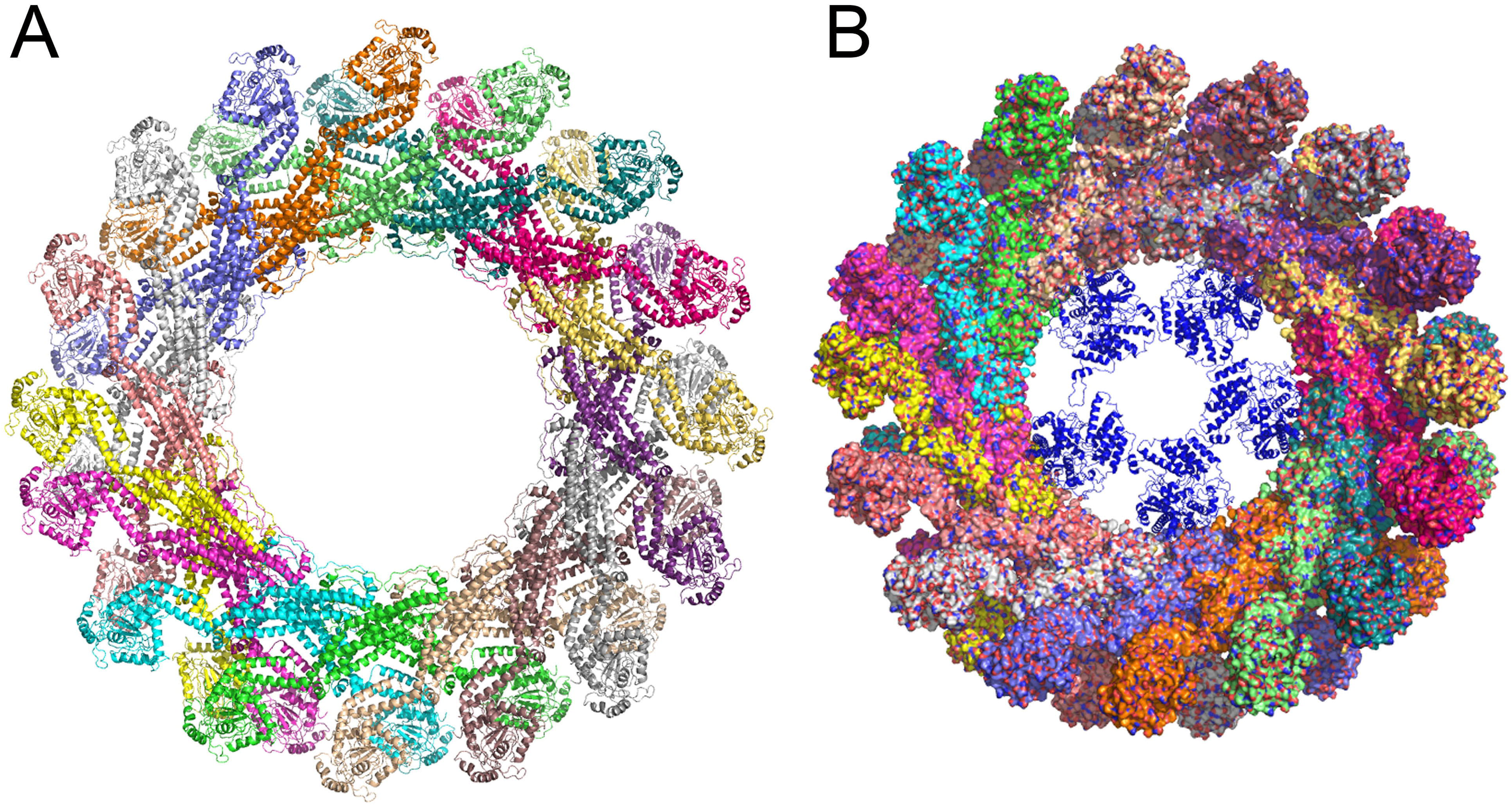
The structural models of NP-polymerase complex, the complete RNP and the RNP-MxA ring. A. The structural model of the NP-polymerase complex. PA, PB1 and PB2 are colored with slate, cyan and yellow, respectively. The two NPs are colored with orange and hotpink, respectively. B. The MDFF-refined NP-polymerase model. D. Detailed atomic interaction between NP and PB1 in the NP-polymerase complex. E. Detailed interaction between NP and MxA in the NP-MxA model.

In this structural model of NP-polymerase complex (Figure 5B), both NPs contact with PA and PB1 subunits of the polymerase. PA contacts both NPs in the model, and so does PB1. PA contact one of the NPs (NP1) by residues in the PA linker, while contact the other NP (NP2) by residues in the PA-arch domain, which is responsible for binding the viral RNA; the PB1 residues that contact NP1 locate in the PB1 fingers domain and that contact NP2 locate in PB1 β-hairpin domain. Some NP residues which localize at the NP-PA or NP-PB1 interface are also the NP interface residues in our NP-MxA dimer structural model (Figure 5C). For example, in one of the NP subunits in the NP-polymerase model, Q42 which interacts with PB1 N58 also interacts with the MxA K556 and S558 in the NP-MxA model; C44 and L49 which interact PB1 K383 also contact MxA D560 and F561 in the NP-MxA dimer model; L47 which interacts with PB1 E387 also interacts with MxA D560; K48 which contacts PB1 S395 also contacts MxA D560; S50 which contacts PA N388 also contacts MxA F561; D51 which forms a hydrogen bond with PB1 K379 and a salt bridge with PB1 386R also contacts with MxA F561 (Figure 5D and 5E). In the other NP subunit in the NP-polymerase model, Y289 which contacts PA P221 contacts MxA D560 and F561 in the NP-MxA dimer model; and R305 which forms a salt bridge contacts MxA D560. Because many NP residues that interact with MxA D560 and F561 residues in the NP-MxA dimer model are responsible for NP binding with PA or PB1 in the RNP complex, MxA may competitively inhibit the binding between NP and PA or PB1 and thus disrupt the assembly of RNP.

## Discussion

The NP and viral ribonucleoprotein (vRNP) complexes of the influenza A virus play a crucial role during the virus infection cycle. During infection, influenza A virus enters the endosome of the host cell by clathrin-mediated endocytosis. After viral membrane fusion occurs in the endosome, viral RNPs are released into the cytosol and then enter the host nucleus by active transport. In the nucleus, the RNPs from the infecting virus serve as templates for the synthesis of viral mRNA as well as anti-genomic, complementary RNAs (cRNA). The cRNAs are replication intermediates that direct the synthesis of nascent virion RNAs (vRNAs). Newly translated NP, PB1, PB2 and PA are imported back to the nucleus and encapsidate the cRNAs and vRNA to form nascent cRNP and RNP structures. The nascent RNPs are then exported from the nucleus with the help of two other influenza virus proteins, M1 and NEP into cytosol. In the cytosol, influenza virus RNPs are transported to the cytoplasmic membrane where they are selectively packaged into budding virions (33) (34).

A current model on the mechanisms of MxA function is that MxA recognizes the incoming vRNPs, then self-assemble into rings, and then arrests RNPs and block its function (22, 35). However, the molecular details of the interaction between MxA and RNPs are unknown. Thus, it is difficult to understand how the planar MxA ring accommodates the helically arranged NPs with sufficient contacts between the L4 loops and the NP. Besides, the incongruous diameters of the MxA ring (24 nm) (22) and vRNP helix filaments (15 nm) (36) also poses a problem for model fitting. We hypothesis the interaction between the MxA ring and the RNP helical filament might be dynamic: all the 32 L4 loops in the MxA ring and the interface residues that we identified in the studies in all NP protomers in the RNP helical filament are potential contact sites and are randomly selected for binding. Binding and unbinding between the MxA ring and the RNP filament occur alternatively, and the binding sites keep changing, but the RNP filament keeps being holding in the MxA ring cavity, and If the interaction model of the vRNP-MxA ring is correct, we envisage the following mechanisms for MxA restriction on the replication of influenza A virus: MxA may capture the vRNP in the cytosol so that the import of the invading vRNPs into the nucleus and the transport of the progeny vRNPs to the budding site are retained. Actually, MxA inhibiting the early stage of influenza A virus infection by preventing transport of the viral genome to the nucleus has been reported (37).

By modelling the NP-polymerase complex of the influenza A virus RNP and analyzing the interface NP-polymerase interface, we show that the interface between NP-polymerase in the modelled NP-polymerase complex partly overlaps with that of NP-MxA. Nevertheless, previous researchers have suggested that MxA targets RNP at the interface of NP-polymerase leading to disruption of the vRNPs and inhibition of influenza virus infection (17).

Our docking analysis also shows that the free NP monomers can form spatially rational complex with MxA-ring. This result casts light on a possible mechanism of MxA: it capture the free newly synthesized NP proteins in cytosol so that their returning to nucleus is retarded and the packing of RNPs is hindered. However, structural and functional data are needed to verify this conclusion.

## Methods

### Modelling NP-MxA complex

Coordinates of MxA (PDB code: 3SZR) and the H5N1 influenza subtype (A/HK/483/97) NP crystal (PDB code: 2Q06) were used for modelling. The missing loops of these two structures were mended by the I-TASSER program (26, 27) The MxA dimer was constructed by aligning two modelled MxA monomers to the MxA dimer obtained by the symmetrical operation on the initial crystal structure of MxA monomer. Subsequently, the resulting MxA dimer was docked onto the NP monomer using the Z-dock program (19), and residues 554-563 in both monomers of MxA dimer was set as the binding sites. The resulting structural models of NP-MxA dimer were further refined by FG-MD program (38).

### Modelling NP-polymerase complex

The structures of PA, PB1 and PB2 of H5N1 influenza subtype (A/HK/483/97) were modelled by I-TASSER program as described above, using the crystal structure of the H17N10 bat influenza A virus (bat/Guatemala/060/2010) polymerase complex (PDB code: 4WSB) (31) as the template. The modelled PA, PB1 and PB2 structure were then align to the template structure (bat polymerase complex) to obtain the modelled polymerase complex of H5N1 influenza A virus. The resulting structural model of the polymerase, together with two NP molecules which make contact with the polymerase, were docked into the electron density map of the RNP-bound polymerase (EMDB entry numbers: 2212) (29), respectively. The resulting structural model of NP-polymerase complex, which contains two NP molecules and the polymerase complex, was further refined by molecular dynamics flexible fitting (MDFF)(32). Briefly, the structural model of NP-polymerase complex was firstly rigid body docked into the electron density map corresponding to NP-polymerase complex extracted from electron density map EMDB-2212 colores program from the Situs package. Then 2 nanosecond MDFF simulation was performed with secondary structure restrains. The scaling factor (ξ) was set to 0.3 and the temperature was set to 300 K.

## Figure legends

Figure S1. The five structural models of MxA dimers. The model shown in (C), which is highlighted in red box, are the model selected for further analysis in the present study.

Figure S2. Modelling the NP-polymerase complex. The structural models of PA (A), PB1 (B), PB2 (C) of H5N1 influenza A virus were generated by I-TASSER program using the corresponding bat influenza A virus polymerase (PDB code: 4WSB as the template). (D). The structural model of the H5N1 influenza A virus polymerase complex was generated by aligning the individual the structural models of PA, PB1, PB2 to the bat polymerase template. (E) Aligning two monomers of the H5 NP and the polymerase structural model of influenza A virus into electron density of RNP-bound polymerase (EMDataBank code: EMD-2212). (F) The atomic structural model of NP-polymerase complex.

Figure S3. MDFF refinement of the NP-polymerase complex. A. The NP-polymerase complex structure model before MDFF refinement. B. The NP-polymerase model after MDFF refinement. C. The RMSD curve for the MDFF simulation. D. The cross-correlation coefficient curve for the MDFF simulation.

